# Network inference reveals novel connections in pathways regulating growth and defense in the yeast salt response

**DOI:** 10.1101/176230

**Authors:** Matthew E. MacGilvray, Evgenia Shishkova, Deborah Chasman, Michael Place, Anthony Gitter, Joshua J. Coon, Audrey P. Gasch

## Abstract

Cells respond to stressful conditions by coordinating a complex, multi-faceted response that spans many levels of physiology. Much of the response is coordinated by changes in protein phosphorylation. Although the regulators of transcriptome changes during stress are well characterized in *Saccharomyces cerevisiae*, the upstream regulatory network controlling protein phosphorylation is less well dissected. Here, we developed a computational approach to infer the signaling network that regulates phosphorylation changes in response to salt stress. The method uses integer linear programming (ILP) to integrate stress-responsive phospho-proteome responses in wild-type and mutant strains, predicted phosphorylation motifs on groups of coregulated peptides, and published protein interaction data. A key advance is that by grouping peptides into submodules before inference, the method can overcome missing protein interactions in published datasets to predict novel, stress-dependent protein interactions and phosphorylation events. The network we inferred predicted new regulatory connections between stress-activated and growth-regulating pathways and suggested mechanisms coordinating metabolism, cell-cycle progression, and growth during stress. We confirmed several network predictions with co-immunoprecipitations coupled with mass-spectrometry protein identification and mutant phospho-proteomic analysis. Results show that the cAMP-phosphodiesterase Pde2 physically interacts with many stress-regulated transcription factors targeted by PKA, and that reduced phosphorylation of those factors during stress requires the Rck2 kinase that we show physically interacts with Pde2. Together, our work shows how a high-quality computational network model can facilitate discovery of new pathway interactions during osmotic stress.

## INTRODUCTION

Cells sense and respond to stressful situations by utilizing complex signaling networks that integrate diverse signals and coordinate what is ultimately a multi-faceted response. In optimal conditions, microbial cells maximize growth at the expense of stress defense by up-regulating growth related processes and suppressing defense strategies. In *Saccharomyces cerevisiae* this is mediated in part by the nutrient sensing RAS/Protein Kinase A (PKA) and TOR pathways [1-3] that promote ribosome production, translation, and proliferation while suppressing activators of the stress response [2, 4]. But upon exposure to severe stress, cells often down-regulate growth-promoting functions while mediating myriad other changes, including in transcription, translation, and post-translational protein modification. Together, these rearrangements produce temporary delay of cell-cycle progression, altered metabolic fluxes, redistribution of the actin cytoskeleton, and other responses. Many of these processes are regulated by post-translational protein phosphorylation; but how cells coordinate many different processes with a single regulatory network during a systematic response remains unclear.

Many studies have characterized phospho-proteomic changes in mutant cells lacking stress-activated regulators. Downstream phosphorylation events requiring those regulators can be readily identified, but most are unlikely to be direct [5]. For example, in the well-studied response to osmotic shock, the HOG pathway is activated to coordinate osmo-induced transcriptome changes [6-8], translation regulation [9-11], cell cycle arrest [12-14], actin reorganization [15-17], and metabolic changes including those that produce intracellular osmolytes [18-20]. However, most of the phosphorylation sites related to these processes do not match the known specificity of Hog1 and are likely controlled by other downstream kinases [21]. Hog1-independent regulators are also activated during osmotic shock [17, 21-23], and other regulators likely remain to be identified. How these connect in a single regulatory network remains unknown.

It is in this context that computational network inference can be particularly powerful. Myriad methods have been developed to analyze phospho-proteomic data; however, many challenges remain. NetworKIN [24] and iGPS [25] leverage known preferences of specific kinases for unique linear sequences around the phosphorylation site, called phospho-motifs [26]. However, these methods work only for kinases with known specificities and they do not place predicted kinase-substrate interactions into hierarchical networks. Other computational frameworks exist for network inference, including methods based on regression [27], Bayesian approaches [28-30], logic-based models [31-35], and ordinary differential equations [36, 37]. These approaches perform best with sufficient biological samples so as to infer statistical dependencies, or when most of the signaling network’s interactions are already known to provide *a priori* guidance. These ideal criteria, however, are not satisfied in typical mass spectrometry-based phospho-proteomic studies, where sample numbers are often limited and the primary goal is pathway discovery unrestricted by prior knowledge. Subnetwork optimization algorithms can be used to extract a small, high-confidence subnetwork from a larger network of interactions, *e.g.* protein-protein interactions (PPI), to explain how signals may propagate through the network [38-41]. Diverse optimization strategies have been used for this task, including methods based on source-target paths [42, 43], network flow [44], and Steiner tree variants [45, 46]. However, these tree-based approaches often cannot handle feedback loops, which are likely very common in signaling networks [40, 47].

To overcome these limitations, we developed an approach to infer the signaling network regulating phospho-proteome changes triggered by stress. We previously developed an integer-linear programming (ILP) method designed to capture the transcriptome-regulating network, by integrating gene-fitness contributions to stress tolerance, wild-type and mutant transcriptomic responses, and phospho-proteomic changes trigged by sodium chloride (NaCl) treatment, which results in osmotic and ionic stress [39]. The resulting network enabled many new predictions about the NaCl response, by identifying new regulators and the regulatory connections between them. However, the inferred network included only 21% of proteins with significant phosphorylation changes during NaCl stress. This is reasonable, since most proteins with phosphorylation changes do not regulate the transcriptome but rather coordinate other aspects of the cellular response [48]. However, it indicates that regulation of much of the cellular stress response is not captured by the previously published network.

Here we adapted our prior ILP approach to infer the NaCl-activated phospho-proteomic regulating network. Our approach first identifies submodules of likely coregulated phospho-peptides that share similar phospho-motifs and mutant dependencies, and then implicates proteins from the PPI network that interact with many of those target peptides. The ILP then assembles those units into a single signaling network, linking upstream regulators we interrogated to downstream phospho-peptide modules dependent on their activity. The method revealed exciting new insights into cellular coordination of disparate physiological responses to stress, several of which we validated through molecular approaches. In particular, the network illuminates cell coordination of cell cycle, metabolism, and growth control during acute stress and points to previously unknown connections between stress-activated and growth-regulating pathways.

## RESULTS

We first profiled the phospho-proteome of wild-type and mutant cells before and after NaCl response. While other studies have interrogated osmo-responsive phospho-proteome changes [17, 21, 39, 49], for optimal network inference we restricted our analysis to data generated in our lab under the identical growth conditions (and used results of other studies as validation metrics of the approach). A full description of the phospho-proteomic data collection is found in the Supplemental Text Section 1. In summary, we used isobaric tagging and mass spectrometry to quantify 8,120 peptides, mapping to 2,049 proteins that were phosphorylated before and/or at 5, 15, or 30 min after treatment with 0.7M NaCl. We leveraged replicates at specific time points to identify 1,249 peptides from 618 proteins that showed reproducible phosphorylation changes (Table S1) in wild type cells responding to NaCl treatment (FDR < 0.05 or meeting our selection criteria, see Methods). These included 479 peptides (38%) with increased phosphorylation and 770 peptides (62%) with decreasing phosphorylation at some time after acute NaCl exposure.

Identifying phospho-peptides dependent on specific regulators enables the ILP to connect those regulators to downstream phosphorylation targets. We therefore also profiled mutants lacking regulators implicated in our previous network, including the osmo-activated Hog1 kinase, the cAMP phosphodiesterase Pde2 that regulates PKA activity [50], and cell-cycle modulating Cdc14 phosphatase. We identified phospho-peptides with reproducible phosphorylation defects upon NaCl treatment (see Methods), implicating 211 defective phospho-peptides in the *hog1*Δ strain and 140 in the *pde2*Δ strain (Table S2). To investigate Cdc14, we used the temperature-sensitive *cdc14-3* mutant paired with an identically treated, isogenic wild type to identify 161 phospho-peptides with in some cases subtle but reproducible phosphorylation defects (see Methods) (Table S2); these were linked to budding, cell polarity and filament formation, GTPase signal transduction, and kinase activity (P < 1×10^−4^, hypergeometric test [51] Table S3), as expected if Cdc14 is inhibited.

### Phospho-proteome network inference

We adapted our prior ILP approach to integrate the wild type and mutant phospho-proteomic changes into a single regulatory network, by linking the three interrogated regulators to their downstream targets defined in the mutant phospho-proteomic analysis. The computational pipeline spans three main steps (Fig 1, complete details in Supplemental Text Section 2):

**Figure 1.**
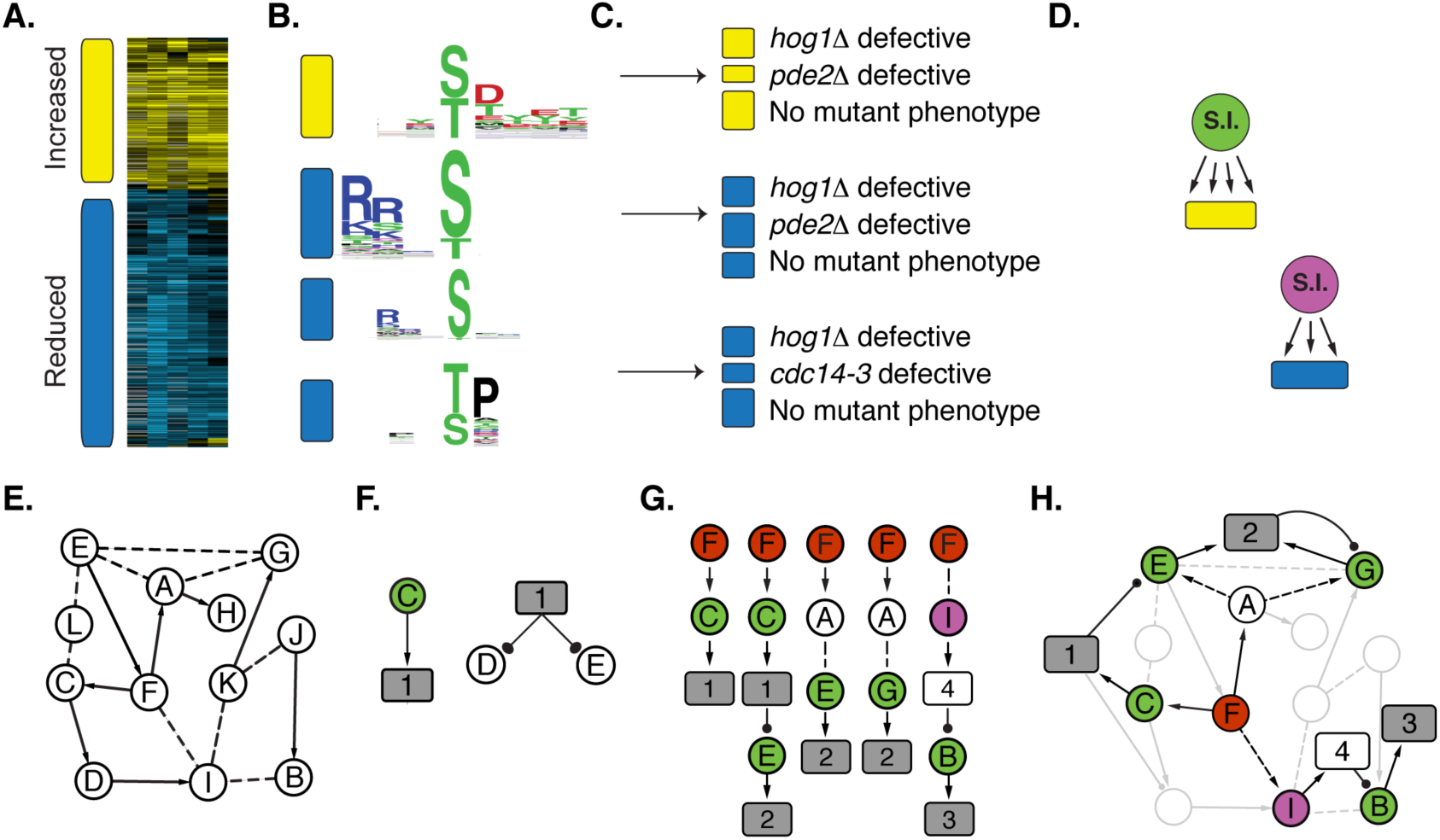
Overview of the inference method. Phospho-peptides are partitioned into submodules based on **(A)** directionality of phosphorylation change, **(B)** shared phospho-motif, and **(C)** mutant dependencies. **(D)** Shared Interactors (SIs) are connected to submodules with a directional edge. **(E)** A background network of previously measured protein-protein (undirected dashed line) and kinase-substrate (directed arrow) is compiled, to which **(F)** directed edges connecting SIs to their identified submodules are added. **(G)** The method then enumerates all paths of a given length from each source regulator (red) to its dependent submodules (grey boxes), traversing through SIs (green, purple) and other proteins in the background network. **(H).** ILP connects the units using a multi-stage objective function, see text for details. Ball-and-stick edges indicate constituent proteins whose peptides are included in a submodule.

#### a) Partition phospho-peptides into potentially co-regulated modules

Many kinases recognize specific phosphorylation motifs and act on suites of targets that harbor those sites [26]. Thus, we identified groups of phospho-peptides that share specific phenotypes, and therefore may be coregulated. First, we grouped peptides into those with increased or decreased phosphorylation after NaCl treatment (Fig 1A). Next, for each group we partitioned peptides based on shared phosphorylation motifs, using the program *motif-X* [52, 53] (Fig 1B). This implicated 17 **modules** of peptides (capturing 71% of NaCl-responsive phospho-peptides) that showed similar directionality in phosphorylation change and shared phosphorylation motifs. Finally, we partitioned each module of peptides based on their dependencies on each of the three interrogated regulators (see Methods). This generated 76 **submodules** of peptides (Fig 1C), including 16 submodules (capturing 159 peptides) that were dependent on Hog1, 19 submodules (100 peptides) dependent on Pde2, 24 submodules (110 peptides) affected by Cdc14, and 17 submodules (563 peptides) with no detectible dependency on any of the three regulators (Table S4). We note that some legitimate targets of the three regulators are likely incorporated in submodules with no nominal mutant phenotype, if the mutant effect fell below our fold-change criteria.

#### b) Implicate potential regulators linked to each submodule

Under the assumption that co-regulated peptides should interact with the same responsible regulator, we leveraged the PPI network to implicate proteins that display more interactions with submodule proteins than expected by chance (FDR < 0.05, hypergeometric test, see Methods) (Fig 1D). We refer to these as **shared interactors** (SIs). A key advantage of this strategy is that it can overcome missing interactions in the published background network, since the SI need not interact with every protein in the submodule (see below). Using this approach, we identified a total of 472 SIs for 54 of the 76 submodules (Table S5). The SIs included 71 kinases, 6 phosphatases, and many proteins of other functional classes - we note that many SIs may not be direct regulators, but instead represent other types of protein interactors (*e.g.* proteins in complex with submodule proteins). SI kinases whose known specificity matched the submodule phosphorylation motif ([54], Kullback-Leibler Divergence, FDR < 0.2%, see Methods) were elevated in confidence as the direct regulator of the submodule peptides.

#### c) Link proteins into a network using ILP

We aimed to connect the SI-submodule units into a signaling network by traversing the background PPI network (Fig 1E). The ILP connects interrogated regulators (Hog1, Pde2, Cdc14, referred to as **sources**) to downstream submodules whose peptides require that regulator for phosphorylation. To do this, we first updated the background network to include edges from each SI to its associated submodules and edges from each submodule to **constituent** proteins whose peptides are included in the submodule (Fig 1F). We then enumerated all possible directed paths of length 3 (discounting submodule-constituent edges) from each source to its dependent submodules (Fig 1G). The ILP then identifies the subnetwork of paths that connect the three source regulators to all downstream submodules, minimizing the number of nodes but maximizing the inclusion of SIs. Many related networks may score equally well; therefore, the output is an ensemble of high-scoring networks with directed, linear paths between sources and submodules (Fig 1H). Because we had few source regulators, we applied a strategy for increasing network diversity within candidate paths [55] by repeating the network inference 1,000 times and each time holding out 5% of proteins from the previous solution. This approach resulted in a richer consensus network compared to our previous approach [39], which combined equivalently optimal solutions to the same ILP. The new approach doubled the number of non-SI proteins included in the network and quadrupled the inclusion of non-SI proteins known to be involved in osmotic stress (as compiled by Chasman *et al* [39, 56-58], see Methods). The resulting ensemble was collapsed into a consensus network, with node and edge confidence values taken as the fraction of ensemble networks in which they were identified. As a final step, we added back submodules that were not included in the consensus but whose SIs were represented. This allowed inclusion of submodules with no detectable mutant dependency, enabling predictions beyond the three source regulators.

The resultant consensus network (at 75% confidence) contained 218 proteins in regulator paths and 51 submodules (encompassing 832 of 1249 phospho-peptides), with 844 edges between them (Fig 2A). The network included 75%, 36.8%, and 62.5% of submodules dependent on Hog1, Pde2, and Cdc14, respectively. The network also predicts the directionality of information flow, suggesting upstream regulatory proteins and downstream targets.

**Figure 2.**
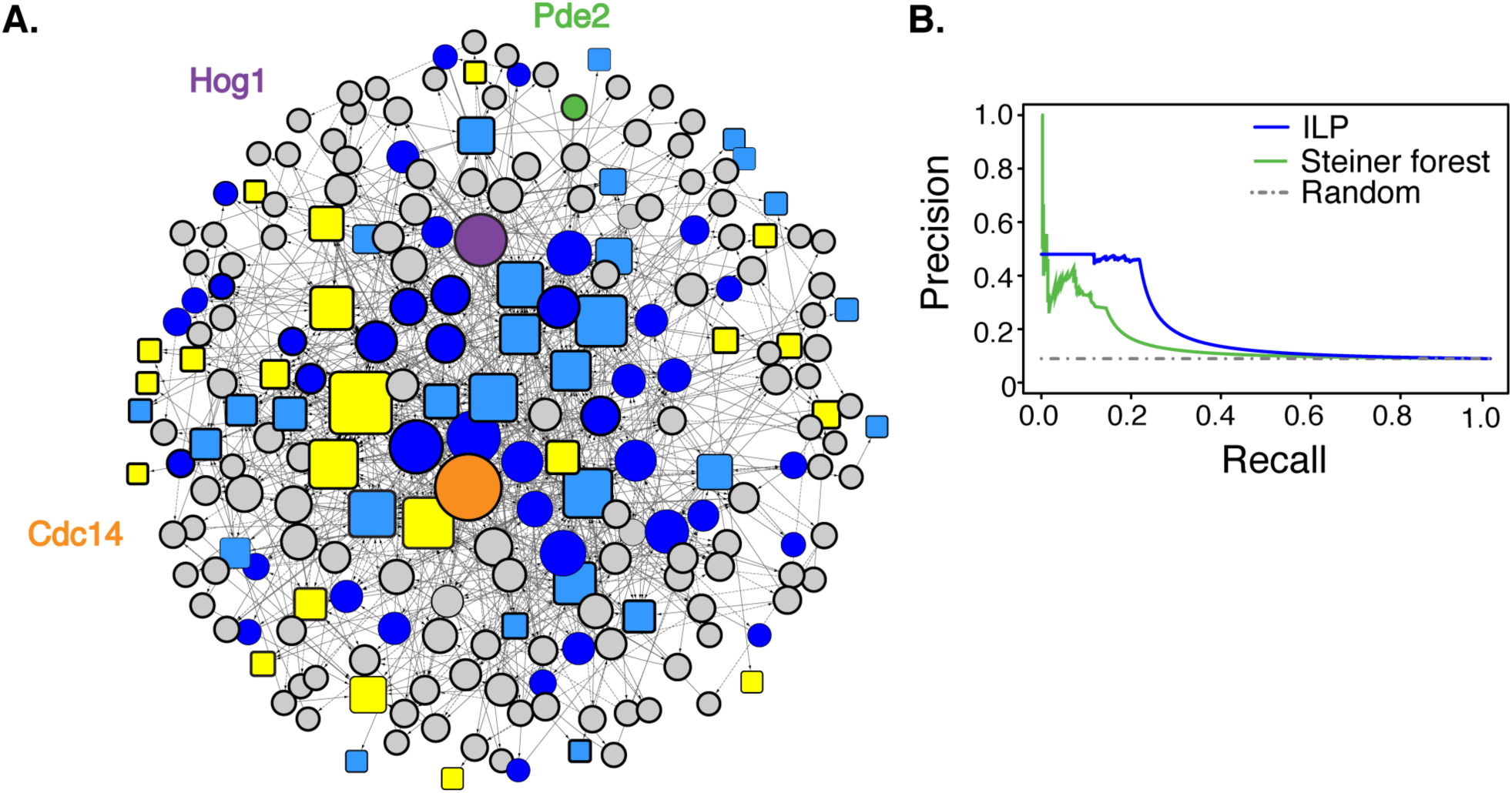
Inferred NaCl-activated phosphorylation signaling network. **(A)** Consensus network at 75% confidence where node size represents degree. Pde2, Hog1, and Cdc14 sources are denoted with green, purple, and orange circles, respectively. Submodules, denoted as rectangles, with increased or decreased phosphorylation in the wild type responding to NaCl are colored in yellow or blue, respectively. (**B)** ILP outperforms an established Steiner forest method [46]. Precision-recall of each method was calculated using a pooled list of true positives, excluding sources, submodules, and shared interactors (see Methods and Supplemental Text Section 3 for evaluation details). *Precision* is the percentage of network proteins that are true positives, while *Recall* is the percentage of true positives retrieved. The AUC was 0.203 for the ILP method and 0.146 for the Steiner forest method (see Methods).

### Computational validation provides strong support for the inferred subnetwork

We assessed the predictive accuracy of the network in several ways. First, the network captured many known regulators of the NaCl response, including proteins in the HOG, RAS/PKA, TOR1, CK2, and Snf1/AMPK pathways. Second, we found that the inferred network was enriched for expected proteins, including kinases (*P* = 3.5×10^−36^, hypergeometric test), proteins that interact with Hog1 or in the HOG pathway (*P* = 8.4×10^−26^) [59-61] and proteins annotated with ‘osmotic’ or ‘stress’ response (*P* = 1.9×10^−14^) [57, 58]. Third, we compared to functional data, leveraging a screen that identified genes important for stress survival after NaCl treatment [62]; indeed, the network is enriched for these functionally important proteins (*P* = 1×10^−3^). Finally, by these metrics the method was significantly better than a random classifier (Fig S1).

Our method also outperformed two other established procedures (see Supplemental Text Section 3 for full details). Compared to NetworKIN, our approach implicated significantly more HOG network components and many more kinases (Supplemental Text, Section 3). Additionally, our approach infers an entire connecting subnetwork, while NetworKIN infers only pairwise interactions. The ILP-based algorithm also outperformed a prize-collecting Steiner forest method for network inference [46]. The Steiner forest method awards prizes for including proteins known to be important and penalizes edges to control the subnetwork size [46]. Compared to the Steiner forest method, our inferred subnetwork captured more relevant genes from multiple true positive lists, scored individually (Fig S2) or pooled together as a single gene set (Fig 2B) (see Methods). It also incorporated more submodules (46 versus 37), and thus captured 463 more phospho-peptides, and identified >3-times more kinases and phosphatases in the final network (Supplemental Text, Section 3). Coupled with biological validations outlined below, this shows that the ILP inference method outperforms existing methods and enables new insights into biology.

### The NaCl-responsive phospho-proteome network captures different functional categories than the previously inferred transcriptome-regulating network

A main motivation was to complement our previously inferred transcriptome-regulating network, so as to more broadly capture cellular signaling and physiological coordination during stress. We expected that, if the inferences are working properly, the two networks should capture proteins involved in different processes, and indeed this is the case (Fig 3). The previously inferred transcriptome-regulating network was heavily enriched for proteins involved in transcription, mRNA transport, chromatin modification, and those localized to the nucleus, among other functions including proteasome degradation (*P* < 1×10^−4^, hypergeometric test [51], Table S6). None of these functions was enriched among the 443 proteins uniquely included (either in regulatory paths or as submodule constituents) in the phospho-proteomic network. Instead, this group was uniquely enriched for proteins involved in endocytosis or found within the eisosome, Golgi apparatus, actin cortical patch, or plasma or vacuole membranes. Annotations related to cell cycle progression, actin cytoskeleton and kinases were enriched among proteins unique to both networks. Beyond these, 145 proteins were identified in both networks and these were enriched for kinases, known osmotic and stress-response regulators, and proteins involved in RAS and PKA signaling, as well as regulators of cell cycle progression/cytokinesis and actin cytoskeleton [51] (*P* < 1×10^−4^) (Fig 3). We propose that many of these represent master regulators coordinating transcriptome changes with other physiological responses to NaCl stress. Indeed, many upstream regulators – including HOG and PKA pathway components – were included in both networks.

**Figure 3.**
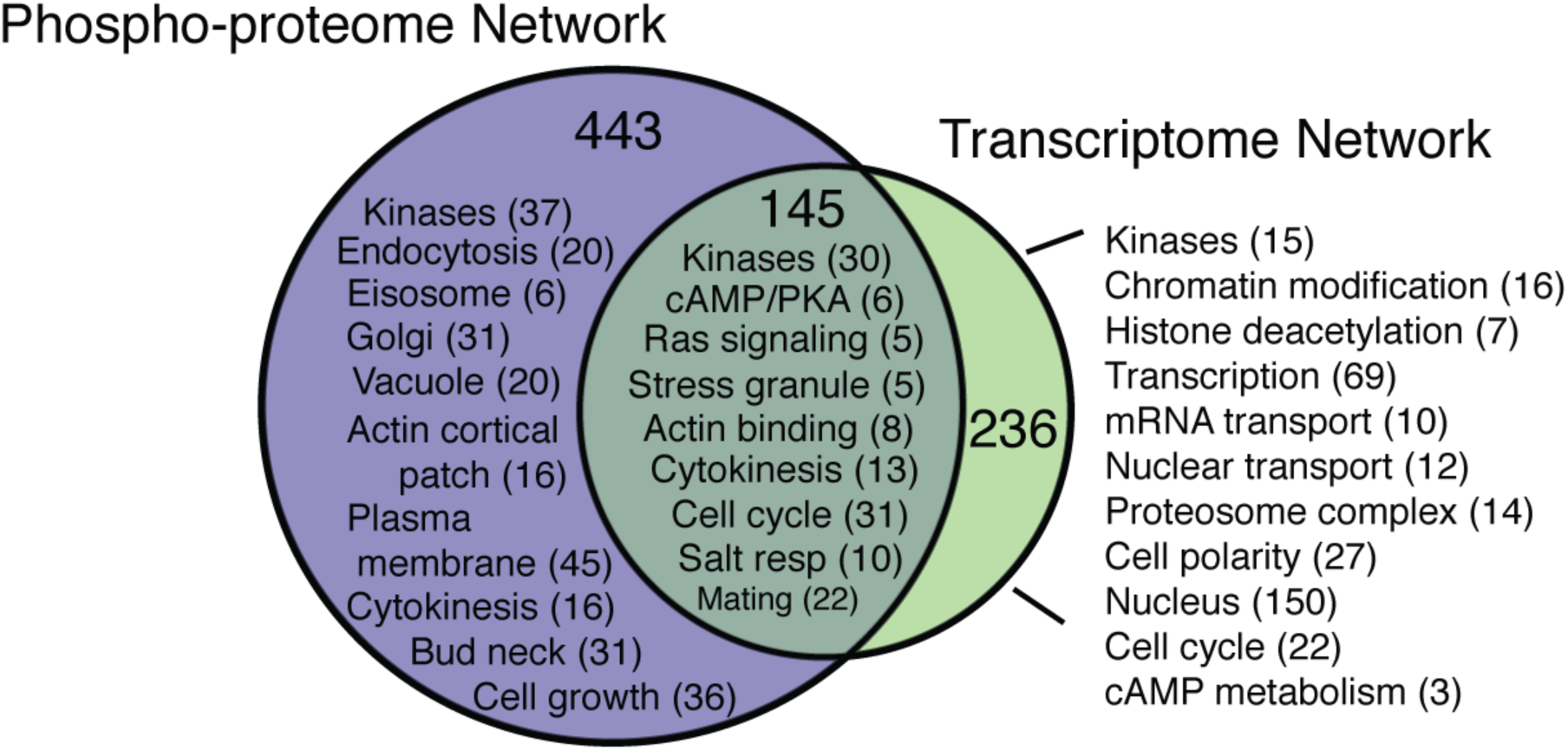
The NaCl-activated networks regulating the phospho-proteome and transcriptome capture unique functional categories. A summary of enriched GO categories (P<1×10^−4^ [51]) for proteins shared and unique to each network. Complete GO enrichments are in Table S6.

### Coordination of cell-cycle control during osmotic shock

Bolstered by the results above, we set out to interrogate the inferred phospho-proteome regulating network, for biological predictions and to provide additional support for the approach. We were interested in leveraging several advantages of our approach compared to other established methods. One key advance is that the method can predict phosphorylation targets by nature of SIs, even when no physical interaction between a predicted regulator and its target has yet to be measured in published datasets. This is important, since interaction datasets are missing many interactions [63], especially those that may be stress specific [64]. Another advantage is that our method can capture feedback loops, which are common in cellular signaling. Feedback is captured when a constituent of a submodule is connected back to that submodule, either as an SI or in a path leading back to that submodule. These provide useful benefits that can expand our understanding of the signaling network.

An example of these benefits is highlighted by the connections between the HOG and cell cycle pathways. Hog1 is known to mediate cell-cycle delay at several phases after osmotic shock. Arrest at G1 is triggered in part by Hog1 phosphorylation of the Cdc28 inhibitor Sic1 [13]. Arrest at G2 is coordinated by a cascade of events, when Hog1-dependent phosphorylation of Hsl1 inhibits its kinase activity, resulting in decreased phosphorylation and thus delocalization of its target Hsl7, which enables accumulation of the Swe1 regulator that phosphorylates and inhibits Cdc28 [12]. Our inferred network captured many of these regulators and information flow between them (Fig 4). For example, the network correctly predicted that Hog1 directly phosphorylates Hsl1 and that Hsl1 is down regulated during NaCl treatment, since all of its connected submodules show decreased phosphorylation. These submodules include Hsl1’s known target, Hsl7, which shows Hog1- and Cdc14-dependent phosphorylation decrease on (Ser-718). While Cdc14 is not a known regulator, it physically interacts with Hsl7 [65], supporting a direct regulatory connection.

**Figure 4.**
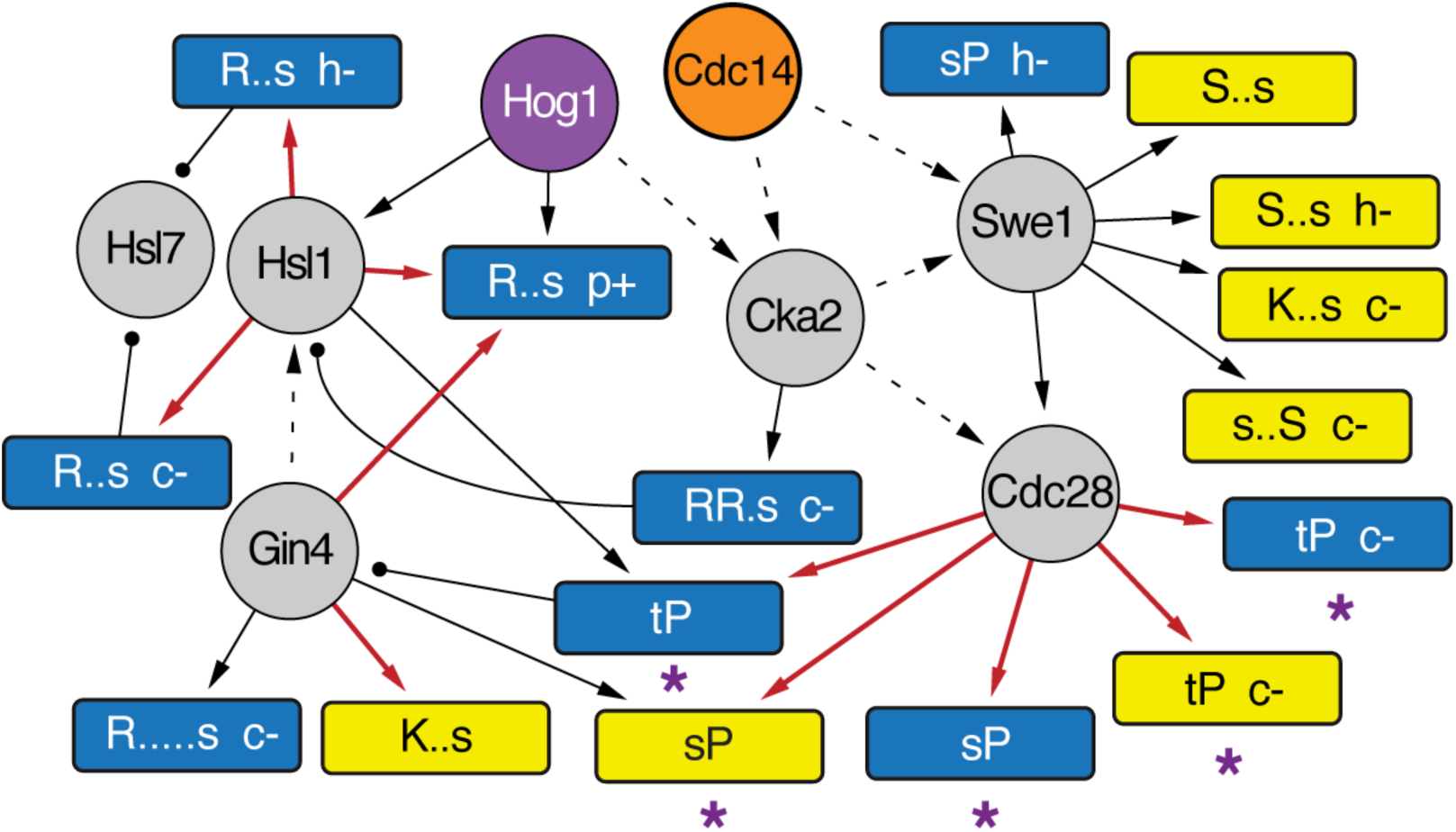
Subnetwork related to cell cycle control. A manually chosen region of the network capturing cell-cycle regulators is shown. Submodules with increased or decreased phosphorylation in the wild type responding to NaCl are colored in yellow or blue, respectively, and annotated by the phospho-motif and mutant phenotype if peptide changes were defective (-) or amplified (+) in the *hog1*Δ (‘h’), *pde2*Δ (‘p’), or *cdc14-3* (‘c’) mutants. Solid arrows represent directed SI-submodule edges or known directional interactions, dashed arrows represent inferred directionality, and ball-and-stick edges indicate protein constituents of the submodule from which the line emits. Red arrows indicate a motif match between the known SI kinase specificity and the target submodule (FDR < 0.2). Asterisks denote submodules containing known Cdc28 target proteins, as curated in Chasman *et al* [60, 61], or phospho-peptides with defective phosphorylation in a strain in which Cdc28 was chemically inhibited [66, 67].

The network also made novel predictions for cell cycle control during NaCl stress. Cdc28 was connected to several submodules whose phospho-motifs match the known Cdc28 specificity, and these spanned 81 phospho-sites with increased phosphorylation and 71 phospho-sites with decreased phosphorylation. Of the combined constituent proteins of all of these submodules, 24% are known Cdc28 targets [60, 61]. Our method predicts that other constituents may represent novel targets. To test this, we compared to two recent phospho-proteomic studies that inhibited *cdc28* analog-sensitive mutants under various conditions [66, 67]. Indeed, the group of predicted Cdc28 target peptides was heavily enriched for sites whose phosphorylation is affected by Cdc28 inhibition (*P* = 5.6 x 10^−15^, hypergeometric test). Only a quarter of these proteins have a known interaction with Cdc28 [60, 61]. Thus, our method elevates the remaining proteins as potential direct targets of Cdc28. Several of these submodules were dependent on the Cdc14 phosphatase, which is known to modulate Cdc28 activity toward different targets [65, 68, 69]. Taken together, these results strongly predict that many of these phospho-peptides are novel direct targets of Cdc28.

### Rck2 is key regulator in the NaCl-activated signaling network

In response to high osmolarity, Hog1 is known to phosphorylate and activate the kinase Rck2, which subsequently phosphorylates targets to decrease translation elongation [11, 70]. We capture a path from Hog1 to a submodule containing Rck2, and then from Rck2 to several submodules whose phospho-motif matches Rck2’s preference and whose phosphorylation is dependent on Hog1 (Fig 5). Only nine percent of the proteins within these submodules are known Rck2 targets [60, 71] leaving the remaining proteins and phospho-sites as novel predictions of our method. We compared these predictions to a recent phospho-proteomics study of an *RCK2* mutant subjected to osmotic stress [21]. Strikingly, phosphorylation of 44% of the novel predicted phosphosites are dependent on Rck2 during osmotic stress (*P* = 2.7×10^−14^, hypergeometric test). One of the constituents is a phospho-peptide mapping to Tps3, a regulatory subunit of the trehalose synthase [72]. While this site on Tps3 (Ser-974) has not been linked to Rck2, Romanov *et al* demonstrated that Rck2 likely phosphorylates several other Tps3 sites [21]. Our results propose that Rck2 also phosphorylates Tps3 on Serine 974.

**Figure 5.**
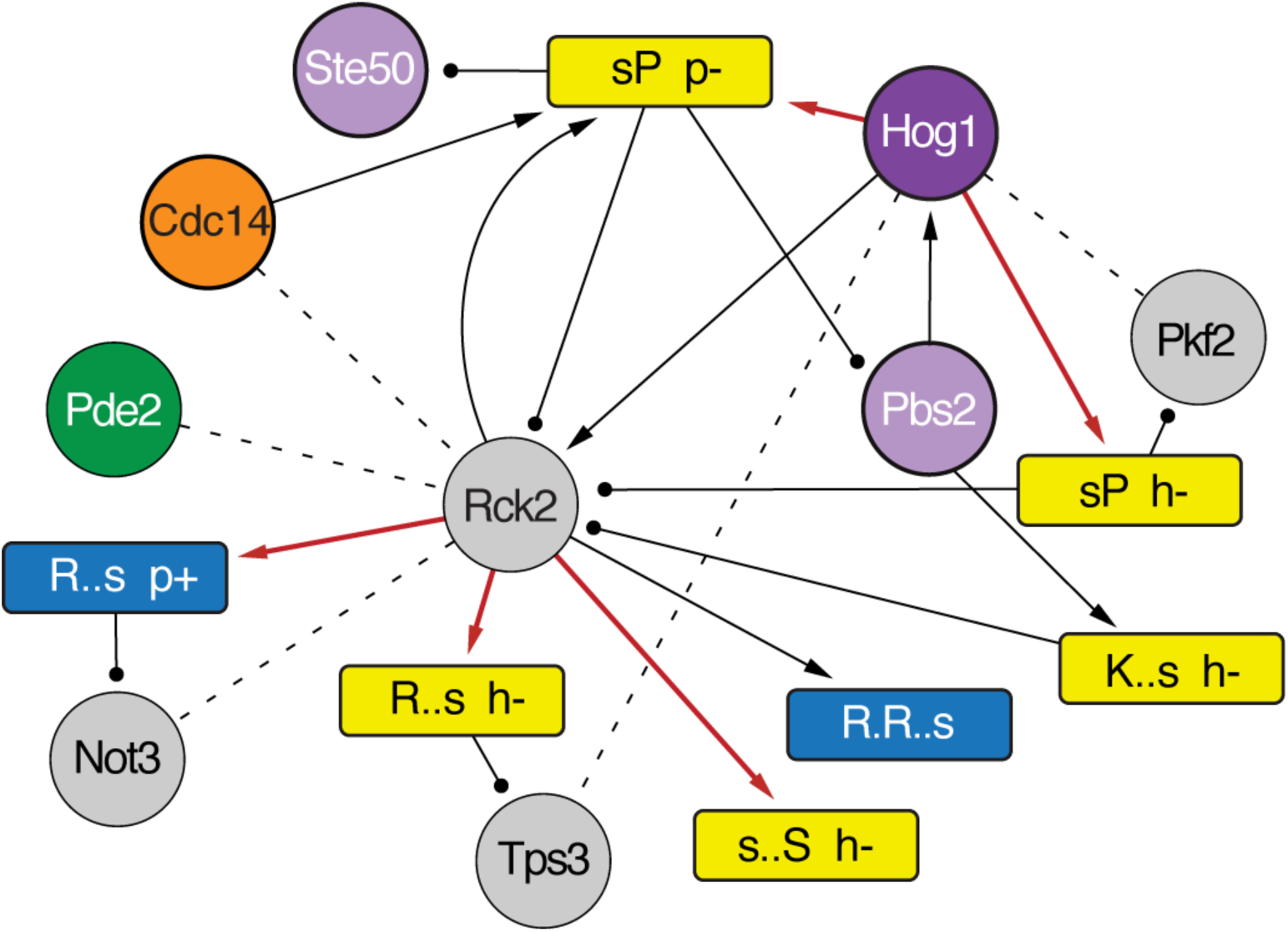
Rck2 is a hub in the osmotic stress-signaling network. A manually chosen section of the network capturing Hog1 and Rck2 regulation is shown, as described in Figure 4. HOG pathway components are colored in purple. Dashed lines represent physical interactions measured by co-IP in this study.

This region of the inferred network captured several other interesting connections. One of the Hog1-connected submodules included Ste50, an upstream regulator known to be phosphorylated by Hog1 as part of a negative-feedback mechanism [73]. Indeed, the submodule captured the known Hog1 target site on Ste50, Ser-202 [73]. Interestingly, NaCl-responsive phosphorylation of the Ste50/Rck2-containing submodule was dependent on Pde2, and in turn one of Rck2’s connected submodules showed a Pde2-dependent phosphorylation change upon NaCl treatment. These results raised the possibility that Rck2 may also be regulated by Pde2 during NaCl stress. Cdc14 was also identified as an SI to the Rck2-containing submodule, although there was no detectible defect in Rck2 phosphorylation in the Cdc14 mutant.

To test these predictions, we immunoprecipitated GFP-tagged Pde2, Cdc14, Rck2, and Hog1 before and 10 min after salt stress and used mass spectrometry to identify other recovered proteins (Table S7). Co-immunoprecipitations (co-IP) validated several of the predicted interactions: IP of both Cdc14 and Pde2 recovered Rck2, whereas IP of Rck2 pulled down Pde2 (albeit just below the threshold used to call co-IPs). Rck2 pull down also recovered Not3, a member of the CCR-NOT complex that was predicted as a novel Rck2 target. That Pde2 is required for Rck2 phosphorylation and predicted downstream Rck2 effects, coupled with the physical interaction between Pde2 and Rck2, strongly suggests that Pde2 is important for Rck2 regulation (see Discussion). IP of Hog1 validated another prediction by recovering the glycolytic enzyme phosphofructokinase Pfk2, which was not previously known to interact with Hog1 but was included in a submodule to which Hog1 was connected and showed Hog1-dependent phosphorylation upon NaCl stress (Fig 5). Together, these results confirm that the ILP network inference can make powerful predictions about regulatory connections.

### New connections and extensive feedback predicted in the Protein Kinase A network

A central player in the osmotic shock response is the PKA pathway, which promotes growth-related processes under optimal conditions but is suppressed to enable defense strategies [2, 74]. How PKA signaling is reduced during stress is not clear. A major portion of our inferred network captured responses linked to PKA (Fig 7). PKA subunits Tpk1, 2, and/or 3 were identified as SIs for 19 submodules, including 6 whose phosphorylation motifs matched the known PKA consensus (R/K-R/K-x-S/T) and an additional 5 matching the lower affinity motif (R-x-x-S) [54, 71, 75]. All of these submodules showed decreased phosphorylation in response to NaCl treatment consistent with reduced PKA activity. The constituent proteins in these submodules are enriched for known PKA targets [60, 61] (*P* = 1.1×10^−5^, hypergeometric test); additional peptides were linked to PKA either through genetic [76] or physical interactions with PKA subunits [77] or defective phosphorylation in PKA catalytic mutants [78].

Interrogating the network revealed new links between PKA signaling and physiology. Collectively, the constituents of PKA-connected submodules with reduced phosphorylation upon NaCl were enriched for proteins involved in budding, cell polarity and filament formation, GTPase- and cAMP-mediated signal transduction, stress response, cell wall structure, and included many kinases (*P* < 1×10^−4^) (Table S8). Fourteen transcription factors were also featured in these submodules (see more below), including stress-responsive factors such as Msn2, Crz1, Sko1, and Dot6 that are inhibited by direct PKA-mediated phosphorylation [79-82]. Of the 17 kinases connected to PKA, over two-thirds are not known to be PKA targets but could represent novel downstream pathways mediated by PKA signaling.

Remarkably, many of the PKA-connected submodules included proteins in the RAS/PKA signaling pathway itself (including Cdc25, Ras2, Cyr1, Ira2, and Tpk3, *P* = 4.8×10^−3^), suggesting extensive feedback control in the PKA pathway (Fig 6). Phosphorylation of several of the captured sites, including Cdc25 Ser-135 and Ras2 Ser-214, has been suggested to play a role in PKA feedback regulation [83-86]. The network also captured phospho-sites of unknown function on the adenylate cyclase Cyr1 (S60), catalytic subunit Tpk3 (S15), and the negative regulator of RAS, Ira2 (S433 and S1018). Phosphorylation of Ira2 (S1018) was previously shown to be decreased in a *TPK3* mutant [78], supporting our supposition of a direct PKA effect. Future studies will be required to dissect the specific effects, but these results suggest significant feedback signaling in the PKA network, which to our knowledge has not been captured previously in inferred regulatory networks.

**Figure 6.**
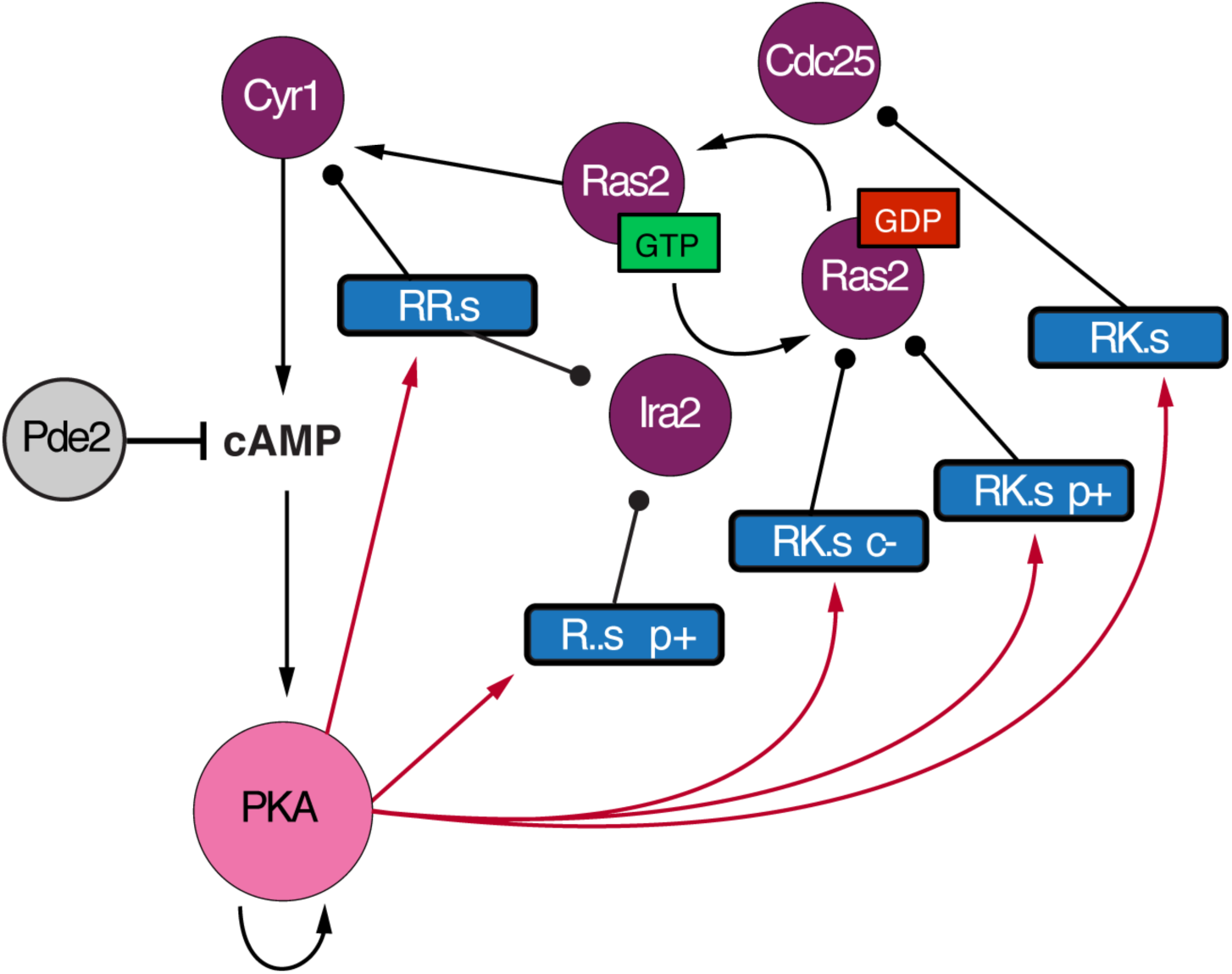
Inferred feedback in the PKA pathway. Shown are all submodules connected to at least one PKA catalytic subunit (‘PKA’) and containing a known regulator in the PKA pathway. The predicted auto-phosphorylation site for PKA in on Tpk3. Figure is as described in Figure 4. Purple/pink circles represent members of the PKA pathway.

### Pde2 interacts with many transcription factors whose phosphorylation is regulated by PKA and Rck2

PKA was connected to many downstream transcription factors, some previously known to be PKA controlled. Several of these, including the general stress factors Msn2/4, Sko1, and Dot6 and calcium-responsive Crz1, lie at the interface of PKA and stress-activated pathways [79, 80, 87, 88]. We noticed that deletion of *PDE2* affected the phosphorylation of many peptides, but relatively few of these were predicted to be directly connected to PKA. Although there is a second phosphodiesterase in yeast (Pde1) that could provide redundancy in PKA regulation, the lack of *pde2*Δ effect on PKA-linked phosphorylation was surprising, since deletion of *PDE2* produces a major defect in salt-responsive transcriptomic changes [39] and causes corresponding defects in acquired stress tolerance after NaCl treatment [62]. But an interesting result emerged from IP of Pde2: nearly a quarter of co-IP’d proteins (17 of 73) were transcription factors, which is much higher than predicted by chance (*P* = 2.6×10^−10^) (Fig 7 + Table S7). Three of these (Msn2, Crz1, and Pho4) are known or predicted PKA targets [71, 79, 80, 87] and another four (Abf2, Rap1, Dig1, Sub1) are part of pathways regulated by PKA [89-93]. Pde2 is critical for normal induction of Msn2/4 targets [39]. This result supports a new model that Pde2 may directly interact with transcription factors to locally regulate cAMP levels (see Discussion).

**Figure 7.**
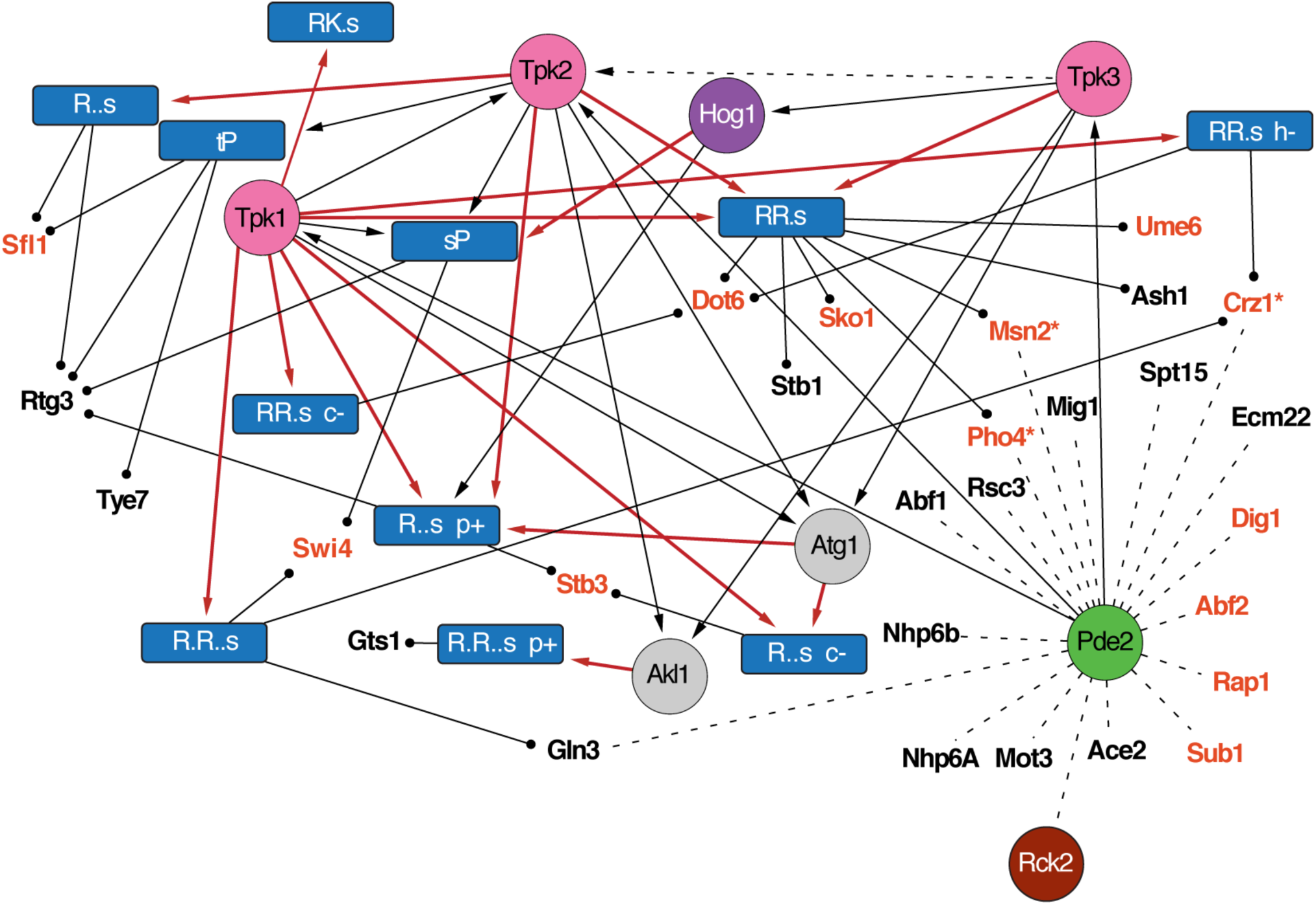
Pde2 interacts with stress-regulated transcription factors. Shown are 10 phospho-repressed submodules (blue rectangles) downstream of at least one PKA subunit and containing at least one transcription factor, as described in Figure 4. Dashed lines without arrows denote Pde2 protein interactions identified by co-IP. Bolded red text denotes factors that are either known or predicted PKA targets [60, 61] or reside in pathways directly regulated by PKA [89-91, 93].

How Pde2 is activated during stress is not known. Our aforementioned finding that Pde2 physically interacts with Rck2 raised the possibility that Rck2 may also influence Pde2 and/or PKA activity. If true, we would expect the *rck2*Δ mutant to have a defect in the salt-dependent decrease in phosphorylation of PKA targets predicted in our network. To investigate, we analyzed the phospho-proteomic response of an *rck2*Δ mutant responding to high NaCl [21]. Strikingly, 35% (118 of 335) of peptides with ≥2-fold higher phosphorylation in the NaCl-treated *rck2*Δ mutant versus wild type harbored the PKA motif (*P* = 4.4×10^−48^). Furthermore, this effect pertained to nearly a quarter of the constituents of PKA-associated, phospho-repressed submodules in our study, more that expected by chance (*P* = 1.8×10^−11^). Many of the affected proteins are known PKA targets, including stress-responsive transcription factors Crz1, Sko1, Yap4, Sfl1, Msn2/4, Maf1, Dot6, and Ifh1 and Rtg1. Other predicted PKA targets were also affected, including Bcy1, Cdc25, Pfk26, and Nth1 that contribute to PKA signaling, growth, or metabolism. Phosphorylation of several of these sites promotes cell growth, including Serine 178 on Pol III repressor Maf1 that alleviates Maf1-dependent repression of rDNA transcription [94]. Taken together, these results suggest that Rck2 plays a role in antagonizing PKA-mediated growth promotion during NaCl stress (see Discussion).

## DISCUSSION

Here we present a novel computational approach to infer the signaling network controlling a dynamic phosphorylation response, in this case the response to the model stressor NaCl. The results not only validate the approach but also highlight the novel insights that can be afforded by network inference.

Our computational method offers several contributions. The ILP’s multi-stage objective function provides flexibility in the inference, allowing the integration of mutant phospho-proteomic data and domain-specific knowledge including SI-submodule connections. It outperformed both NetworKIN and the Steiner forest method implemented here by several metrics. The approach to link SIs to submodules affords several other opportunities. First, it enables the prediction of novel protein interactions that are currently missing from the protein interaction network. While these remain predictions until proven by other methods, they present new hypotheses for subsequent direct testing. In spite of the high false-negative rate of Co-IP analysis via mass spectrometry [95-97], we successfully validated several new predicted interactions (Fig 5). This included interactions between Rck2, Cdc14, Hog1, and several predicted targets (Table S7). Another advantage is that SI-submodule relationships predict not only the targeted protein but also the specific phospho-site modified by that regulator. We showed that predicted target sites were indeed defective in cells lacking functional Cdc28, Rck2, or PKA. Our capacity to predict both the upstream regulator and the targeted site(s) presents an exciting opportunity to test the specific effects of those sites, through phospho-mimicking mutations [98]. Finally, the ability of our approach to predict signaling feedback will be particularly useful in dissecting how cellular signaling is amplified, attenuated, and augmented by pathway activation.

Despite these advances, there are several limitations of our approach as currently implemented. One challenge is in how peptides are partitioned into likely co-regulated sites. On the one hand, co-regulated targets may be inappropriately split into multiple submodules due to overfitting. For example, PKA can phosphorylate both RKxS and RRxS sequences [71], yet *motif-X* partitions peptides containing these motifs into separate modules. Such overfitting may hinder the subsequent identification of SIs due to small numbers of constituents in each submodule. On the other hand, sites recognized by different kinases can in theory be incorporated into the same submodules, if those kinases share the same specificity and mutant dependencies. Proline-directed Hog1 and Cdc28 both recognize SP motifs while PKA and Sch9 regulate basophilic sequences [54]. This is a key consideration when formulating hypotheses for subsequent testing.

Nonetheless, the inferred network proved powerful in providing insights into stress biology, including the interplay between stress defense and growth-promoting pathways. PKA regulators were prominently featured in the network, suggesting myriad new PKA targets involved in cytoskeletal rearrangements, growth control, feedback signaling and stress-dependent transcription. How PKA is down-regulated during the NaCl response remains unclear. Deletion of *PDE2*, one of the two cAMP phosphodiesterases in yeast, did not produce a major defect in predicted PKA-dependent phosphorylation events despite a major defect in the NaCl-responsive transcriptomic changes [39]. In mammalian systems, phosphodiesterases have been implicated in creating micro-environments of low cAMP for local PKA control [99-101], reviewed in [102, 103]. One possibility is that Pde2 locally regulates cAMP surrounding transcription factors, *e.g.* in the nucleus or on the target-gene promoters, rendering a major defect in gene expression but little apparent effect on bulk cellular phosphorylation profiles. Future experiments will be required to test this model. Our analysis also suggests that Rck2 plays an important role in PKA regulation. Data from Romanov *et al.* show that the *rck2*Δ mutant treated with NaCl shows significantly higher phosphorylation of many PKA targets compared to a NaCl-treated wild type [21]. Interestingly, *hog1*Δ mutants (in [21] and here) do not produce the same striking effect, suggesting Hog1 independence. *RCK2* mutants share several phenotypes with negative regulators of PKA: cell lacking *RCK2*, *PDE2*, and the negative RAS/PKA regulator *IRA2* are sensitive to osmotic shock, while *rck2*Δ and *ira2*Δ mutants are resistant to the phosphodiesterase inhibitor caffeine (albeit detected in different studies [104, 105]). Intriguingly, Rck2 has sequence and functional homology with mammalian MAPKAP2, a stress activated kinase activated by the Hog1 ortholog p38 [106]. MAPKAP2 was recently shown to directly regulate Pde4 in human cells [107], supporting our conjecture that Rck2 regulates Pde2 and/or PKA in yeast as well.

A major challenge going forward in phospho-proteomic research is dissecting functional consequence of phosphorylation events. While some phosphorylation events produce major consequences in the cell, many likely fine tune dynamics, noise, and memory [108] and others may be inconsequential [17, 109]. Functional dissection of the outcome of such events (*e.g.* using phospho-mimicking mutations at specific sites) will be an important step in understanding cellular coordination during stress.

## MATERIALS AND METHODS

### Strains and growth conditions

Strains used were of the BY4741 background unless otherwise noted [110] (Thermo Scientific, Waltham, MA). The *cdc14-3-URA3MX* mutant was kindly provided by Michael Tyers [111]. GFP-tagged strains were obtained from the yeast GFP collection [112] (Thermo Scientific, Waltham, MA). Correct knockouts or GFP integrations were verified by diagnostic PCR. Wild-type samples for phospho-proteomics were grown for at least 7 generations in YPD at 30°C to log phase, followed by collections before and 5 min (in biological triplicate), 15 and 30 min (in one replicate time course) after treatment with 0.7M NaCl. The *hog1*Δ and *pde2*Δ mutants, along with a paired wild type, were collected before and 5 min after NaCl treatment in biological duplicate. The *cdc14-3-URA3MX* strain and its parent, MT1901, were grown for 7 generations at 25°C followed by centrifugation for 2 min at 3K, decanting, and resuspension of cells in pre-warmed media at the non-permissible temperature (37°C). After 2 hr at the non-permissible temperature, cells were harvested before and 5 min after 0.7M NaCl treatment. Cdc14 inactivation was verified by a high fraction of dumbbell shaped cells [68]. Cells were collected by brief centrifugation, flash frozen, and stored at -80C.

### co-Immunoprecipitation (co-IP) assays

Hog1-GFP, Rck2-GFP, Cdc14-GFP, and Pde2-GFP were grown for at least 7 generations in YPD and harvested before and 10 min after 0.7M NaCl treatment, followed by immediate flash-freezing. BY4741 was included as an untagged control. Yeast cells were resuspended in lysis buffer (50 mM HEPES-KOH pH 7.5, 140 mM NaCl, 1 mM EDTA, 0.5% NP40, and 0.1% Na-Deoxycholate with protease inhibitors (Millipore, Billerica, MA) and phosphatase inhibitor NaF (Thermo Scientific, Waltham, MA)), and lysed by glass bead milling (Retsch, Newton, PA). Immunoprecipitations were performed by incubating ∼12.5 mg of protein lysate with 25 μL of GFP-Trap MA beads (Chromotek, Germany) for 1.5 hr at 4°C. Beads were washed twice with 1 mL wash buffer (50 mM HEPES-KOH pH 7.5, 140 mM NaCl, 1 mM EDTA, 0.5% NP40), followed by 2 washes with 1 mL Tris wash buffer (150 mM NaCl, 10 mM Tris-CL pH 7.5, and 0.5 mM EDTA). Proteins were eluted with 20 μl 0.5% formic acid and lyophilized in a speed vacuum. Each mutant was interrogated in biological duplicate with a matched un-tagged wild-type strain as a control.

### Mass spectrometry summary

A brief overview of mass spectrometric analysis is provided here, with additional details in the supplement. Cell pellets were thawed on ice, washed twice with 1 ml ice-cold water, and resuspended in lysis buffer (8 M urea, 50 mM Tris pH 8.0, and protease and phosphatase inhibitor cocktail table, Roche, Indianapolis, IN) and rigorously vortexed. Yeast cells were lysed by glass bead milling (Retsch, Newton, PA). Briefly, 500 μl of acid washed glass beads were combined with 500 μl of resuspended yeast cells in a 2 ml Eppendorf tube and shaken at 4°C 8 times at 30 hz for 4 min with a 1 min rest in between. Bradford Protein Assay (Bio-Rad, Hercules, CA) was used to measure final protein concentration. Proteins were reduced by incubation for 45 min at 55 °C with 5 mM dithiothreitol. The mixture was cooled to room temperature and alkylated by addition of 15mM iodoacetamide in the dark for 45 min. The alkylation reaction was quenched with equivalent amount of 5 mM dithiothreitol.

Cell lysates were prepared and diluted with 50 mM Tris to a final urea concentration of ∼ 1.5 M before the addition of trypsin in 1:50 ratio (enzyme:protein; Promega, Madison, WI). Mixtures were incubated overnight at an ambient temperature, acidified by the addition of 10% TFA, desalted over a Sep-Pak (Waters, Milford, MA), and lyophilized to dryness in a SpeedVac (Thermo Fisher, Waltham, MA). Peptides were labeled with tandem mass tags (Pierce TMT, Rockford, IL), according to the manufacturer’s instruction. Labelled peptides were then mixed in 1:1 ratio, and the resulting mixture was desalted over a Sep-Pak. ∼2.5 mg of the labelled peptide mixture were used to enrich for phospho-peptides via immobilized metal affinity chromatography (IMAC) using magnetic beads (Qiagen, Valencia, CA), according to the published method [113]. High pH reverse phase liquid chromatography was used to fractionate enriched phospho-peptides. Peptides were analyzed on a quadrupole-ion trap-Orbitrap hybrid Fusion^®^ or Fusion Lumos^®^ mass spectrometer (Thermo Scientific, San Jose, CA), as described in detail in the Supplemental Text.

### Mass-spec data analysis

The raw data corresponding to TMT-labelled peptides were searched against *Saccharomyces* genome database (SGD) of yeast protein isoforms (downloaded 12.02.2014) and processed using the COMPASS software suite [114], with FDR correction at the peptide and protein level (<1%). TMT reporter region quantification was performed using an in-house software TagQuant, as previously described [115]. Briefly, the raw reporter ion intensity in each TMT channel was corrected for isotope impurities, as specified by the manufacturer for the used product lot, and normalized for mixing differences by equalizing the total signal in each channel. In cases where no signal was detected in a channel, the missing value was assigned with the noise level of the original spectrum (i.e. noise-band capping of missing channels), and the resultant intensity was not corrected for impurities or normalized for uneven mixing. The raw data corresponding to the Co-IP analyses were processed using MaxQuant (Version 1.5.2.8; [116]). Searches were performed against a target-decoy database of reviewed yeast proteins plus isoforms (Uniprot, downloaded January 20, 2013) using the Andromeda search algorithm with precursor search tolerance of 4.5 ppm and a product mass tolerance of 20 ppm, as further described in Supplemental Text (peptide and protein FDR < 1%). Proteins were identified by at least one peptide (razor + unique) and quantified using MaxLFQ with an LFQ minimum ratio count of 2. The ‘match between runs’ feature utilized, and MS/MS spectra were not required for LFQ comparisons.

To implicate co-IP’d interactors from contaminating proteins, we selected proteins that were i) identified in both biological IP replicates (either before or 10 min after NaCl treatment), and ii) were ≥2-fold more abundant in the GFP-tagged pulldown compared to the untagged control. Protein-protein interactions were subsequently classified as salt-dependent or salt-independent by calculating the log_2_ ratio of protein abundance at 10 min versus pre-stress (for this calculation, missing data was imputed as the lowest 1% of intensity scores from that run). Proteins with ≥1.5-fold differential abundance before versus after NaCl in both replicates were classified as salt-dependent interactions.

### Functional enrichments and lists of true positives

Unless otherwise noted, functional enrichments were calculated by the hypergeometric test using FunSpec [51], taking categories with P<1×10^−4^ (representing the Bonferroni-corrected threshold) as significant. Several true positive lists were used to assess network accuracy: ‘Osmotic stress response genes’ contained 110 previously curated [39] proteins that are either HOG pathway components [56, 58], contain ‘osmotic’ or ‘osmolarity’ in their *Saccharomyces* Genome Database (SGD) annotation [57], or are annotated as ‘stress regulator’ and identified in at least one publication as osmotic stress associated. The ‘HOG pathway’ list contained 78 proteins that either physically interact with Hog1 in the background network, as compiled by Chasman *et al* [39], or were identified as HOG pathway components in other studies [56, 59]. The ‘Kinase-phosphatase’ list contained 129 kinases and 30 phosphatases in yeast [111]. The ‘Shared Interactor’ list contained 472 proteins that were identified as SIs in this study.

## COMPUTATIONAL PIPELINE

### Defining phospho-peptide submodules

To identify changing phospho-peptides with significant phosphorylation change in replicate time points, we input count-based phospho-peptide intensities to edgeR [117] and took (FDR <0.05) as significant. In addition, we selected proteins with ≥1.5-fold change in 2 of the 3 paired replicates comparing to samples from paired, untreated cells. We added to this phospho-peptides from the single time course that had at least a 1.5X change in both the 15 and 30 min time points, or a single instance of ≥2X at the later time points. This identified 1,249 phospho-peptides that respond to NaCl, which were split into peptides that increased or decreased phosphorylation (Table S1). Further partitioning based on hierarchical clustering of temporal profiles did not provide any benefits in identifying other clusters (not shown).

To identify reoccurring motifs, *motif-X* [52] was run separately on phospho-peptides with NaCl-dependent increases or decreases, using the following parameters: extend from: SGD yeast proteome; central character: s* or t*; width: 13; occurrences: 10; significance: 1×10^−6^. This yielded 17 motifs that capture 80% of all changing phospho-peptides. Groups were further split based on mutant dependencies as follows: for *hog1*Δ and *pde2*Δ mutants, peptides with ≥1.3-fold difference in abundance in both replicates (or 1.5-fold for peptides detected in only one of the two experiments) compared to wild type were considered affected in the mutant. Due to the well-known phenomenon of inference-induced ratio compression associated with TMT quantification, all measured changes were likely underestimated [118]; we therefore determined the aforementioned cutoffs manually so as to capture known targets of the regulators. Functional enrichments of the selected proteins support the approach: peptides identified as Hog1 dependent were enriched for ER to Golgi transport and cell growth, whereas Pde2 was enriched for cell growth, actin cytoskeleton, budding, cell polarity and filament formation, and guanine nucleotide exchange factors (GEFs) (Table S2) (P < 1×10^−4^) [51]. Twenty-three percent of the proteins with Hog1-dependencies and found in the enriched groups require Hog1 for NaCl-responsive phosphorylation changes [21], validating our selection criteria. Fold-changes smaller in the mutant than the wild type were termed **defective**, changes that were greater in the mutant than wild type were termed **amplified**, and phospho-peptides that failed to meet these criteria were classified as having **no-phenotype** in the mutant. We relaxed the fold-change cutoffs for the *cdc14-3-URA3MX* mutant. This mutant exhibited smaller but still reproducible defects in phosphorylation compared to the other mutants, including for known Cdc14 targets and proteins linked to the cell cycle. The subtle but reproducible effects may be because Cdc14 functions in only a subset of cells in the population, other phosphatases act partially redundantly, or the required experimental procedure limited our power to detect defects. We therefore used relaxed criteria to identify phospho-peptides affected by Cdc14, requiring a reproducible 1.15-fold defect in phosphorylation compared to the mock-treated wild type, or a 1.3-fold defect for peptides detected in a single sample. Phospho-peptides identified as Cdc14-affected were enriched for functions related to Cdc14 activity, including cell cycle, budding, cell polarity and filament formation, in addition to GTPase signal transduction and kinase activity (Table S2). Together, these criteria were used to subdivide each of the 17 modules into 76 submodules. Peptides affected by multiple mutants were represented in each of the corresponding submodules, rather than placed into submodules with multi-mutant effects, due to small numbers.

### Identification of Shared Interactors and phospho-motif matches

We used a background PPI network as compiled by Chasman *et al* [39], which included protein-protein and kinase-substrate interactions measured in multiple high-throughput studies (or individual low-throughput studies) compiled from various databases [60, 61, 111, 119, 120]. For each submodule of peptides, we identified proteins from the PPI network that showed more interactions with submodule constituent proteins than expected by chance (hypergeometric test, FDR < 0.05), which yielded 472 SIs linked to one or more of 54 submodules. Edge directionality was determined based on known kinase-substrate relationships: **input** edges from the SI to the submodule were assigned for SIs with at least one directed interaction with submodule constituents or for SIs with known, non-directional interactions with those constituents. We classified SIs with non-directional interactions as inputs because these interactions might be directed, but were not identified as such because of previous experimental design limitations or tested conditions. In contrast, directed edges from the submodule to the SI were defined as **output** edges if all of interactions were directed toward the SI; these edge directionalities were considered in the subsequent ILP.

For final network visualization, we categorized edges as **motif-match** if the known specificity of its SI kinase matched the submodule motif identified by *motif-X*. We first generated a position weight matrix (PWM) for module peptides distinguished by *motif-X*, adding a pseudocount for each amino acid to prevent overfitting (Table S9). These were compared to protein-array defined specificities for 63 kinases from Mok *et al* [54], containing background-corrected phosphorylation signal intensities that were normalized to total signal intensities for all amino acids. We converted to frequencies by summing signal intensities for all amino acids at a position (after addition of a pseudocount) and dividing by the summed intensities. Adding a pseudocount was necessary as some amino acids were not detected in the protein-array, and Kullback-Leibler Divergence (KLD) requires non-null values at all positions. Next, we compared the module PWMs to the adjusted Mok PWMs using Kullback-Leibler distance. To assess statistical significance of the matches, we generated a randomized distribution of KLD scores by permuting within-column values, and then shuffling the columns for each kinase PWM from [54] 1000 times, generating 63,000 KLD scores per module. FDR was taken as the number of randomized KLD scores that had a smaller KLD distance than the observed value.

A motif-match designation was assigned to kinases with the smallest FDR scores that also belong to a kinase group (as assigned by Mok *et al*) that is known to recognize the module motif. Kinases that were not in the Mok *et al* dataset were classified as having **unknown-recognition motif** relationships to all modules.

### Integer linear program method for subnetwork inference

We use an integer linear program (ILP) to select paths through the modified background network to link interrogated source regulators to their downstream phospho-peptide submodules, minimizing the number of intermediate nodes used by all paths while maximizing the inclusion of shared interactors (SIs). The ILP is a modified version of what was proposed by Chasman *et al* [39]. An overview of the approach is described below with a detailed description provided in Supplemental Text Section 2.

We first augmented the background network with 1,835 edges capturing SI-to-submodule *input* edges and submodule-to-constituent or submodule-to-SI *output* edges as described above (Table S5). We next enumerated all possible acyclic paths of up to 3 edges (discounting submodule-to-constituent edges), between interrogated *source* regulators (Hog1, Pde2, Cdc14) and the phospho-peptide submodules that exhibit a phenotype, executed as a depth-first search through the background network. Submodules without any mutant dependencies were included as nodes in the background network and may appear as intermediates in paths but not as termini. We assign a binary variable to each network element (node, edge, and path) to represent whether the element is selected for inclusion in the subnetwork or not. For undirected edges, we also assign a directionality variable, *d* which is set to 1 if the edge is selected in the ‘forward’ direction (determined by lexicographic order of the node names) and 0 otherwise. We also make use of a variable *c*_*s,m*_ for each source-submodule pair that indicates whether the pair has been connected by a selected path.

We developed a multi-part objective procedure similar to our previous work [39] and described in detail in Supplemental Text Section 2. At each step, the result of the solution is added to the IP as an additional constraint that must be satisfied during the next optimization step: 1. Maximize the number of source-submodule pairs connected by a selected path. 2. Find the maximum number of SI edges that can be included in a valid subnetwork. 3. Minimize the number of intermediate nodes that are not *sources*, *submodules*, or SIs. 4. Maximize the number of selected paths. We implemented a step to increase diversity in the final solutions based on a previous approach [55]. Specifically, after each solution we randomly hold aside (that is, fix its *y* variable to 0) with 5% probability all non-source, non-submodule nodes from the previous solution. As a result, about 5% of previously selected nodes are held aside in each iteration. We repeated this procedure 1000 times. After pooling these solutions, we defined a consensus network based on source-submodule paths found in at least 75% of all solutions. As a final step, we added to this consensus all submodules without mutant phenotypes whose SI was included in the consensus, with a direct edge emerging from its SI to that submodule. In all, the final network included 218 proteins in regulator paths and 51 submodules (together capturing 832 of 1249 phospho-peptides), with 844 edges between them. Precision-recall analysis is described in Supplemental Text Section 3.

### Data Availability

All raw proteomics data files were deposited into PRIDE repository (Accession #PXD006192). Reviewers can use the following information to download files during the peer review process:

Username

Password: jBbJVRcs

### Computational Pipeline Availability

All scripts are available at the following GitHub repository under the GPLv3 license: https://github.com/mmacgilvray18/Phospho_Network

## ACKNOWLEDGEMENTS

This work was supported by NIH R01GM083989 to APG, P41 GM108538 and R35 GM118110 to JJC, and NSF 1553206 to AJG. MEM was supported by an NIH training grant (T32 HG002760); MP was supported by the Department of Energy funded Great Lakes Bioenergy Research Center (BER DE-FC02-07ER64494). We thank Michael Ferris for computational resources.

